# Significant improvement of reprogramming efficiency by transient overexpression of ZGA inducer Dux but not Dppa2/4

**DOI:** 10.1101/803593

**Authors:** Lei Yang, Xuefei Liu, Lishuang Song, Anqi Di, Guanghua Su, Chunling Bai, Zhuying Wei, Guangpeng Li

## Abstract

Cloned animal have been reported by somatic cell nuclear transfer (SCNT) for many years. However, the SCNT is extremely inefficient, and zygotic genome activation (ZGA) is required for SCNT and chemical mediated reprogramming. To identify candidate factors that facilitate ZGA in reprogramming, we performed siRNA-repressor and mRNA-inducer screening, which revealed Dux, Dppa2, and Dppa4 as key factors enhancing the ZGA in SCNT. Direct injection of ZGA-inducers had no significant effect on the SCNT blastocyst formation, and even destroyed the ZGA. Through a inducible Dux transgenic mouse model, we demonstrate that transient overexpression of Dux not only improved SCNT efficiency, but also increased the efficiency of chemical reprogramming. Transcriptome profiling revealed that Dux treated SCNT embryos were similar to fertilized embryos. Furthermore, transient overexpression of Dux combined with inactivation of DNA methyltransferases (Dnmts) further promote the overall development of SCNT-derived animals. These findings enhance our understanding of ZGA-regulators in somatic reprogramming.

## Introduction

The scientists at the Institute of Neuroscience (Chinese Academy of Sciences), recently cloned the world’s first monkeys Zhong Zhong and Hua Hua (the Zhong Hua is a mandarin term for china) (Z. Liu et al., 2018). After the first mammal Dolly sheep to be cloned (Wilmut, Schnieke, McWhir, Kind, & Campbell, 1997), the created non-human primate has once again attracted worldwide attention. Indeed, successful cloning of the sheep and monkey not only made reproductive cloning possible, but also raised the possibility of therapeutic cloning. Somatic cell nuclear transfer (SCNT) is the technical term for cloning in which terminal differentiated somatic cell is transferred into enucleated oocyte, in which the cytoplasmic factors reprogram the nucleus to become a zygote-like state (Matoba & Zhang, 2018). Both SCNT derived and Yamanaka factors induced pluripotent stem (iPS) cells have the potential to generate patient-specific pluripotent stem cells for replacement therapy. Compared with iPS, the SCNT reprogramming is oncogene-free methodologies. Thus, SCNT derived pluripotent stem cells would be more suitable for human therapeutic applications.

Following fertilization, the newly formed zygotic genome is activated through a process known as the zygotic genome activation (ZGA), which enables zygote can subsequently development to the adult animal (Lu & Zhang, 2015). A similar mechanism is likely used in ZGA of SCNT embryos (T. Gao et al., 2007; Matoba & Zhang, 2018). Indeed, our and others recent results have revealed that incomplete ZGA is a major reprogramming barrier (W. Liu et al., 2016; Matoba et al., 2014; Wang et al., 2018; Yang, Song, Liu, Bai, & Li, 2018). Although many advances have been made in SCNT technology via different epigenetic regulators, the SCNT for producing cloned animals still inefficient.

The transcription factor of double homeobox (Dux) was identified as a key inducer of ZGA in normal fertilized embryos (De Iaco et al., 2017; Hendrickson et al., 2017; Whiddon, Langford, Wong, Zhong, & Tapscott, 2017). Very recently, two independent studies found that transcription factors developmental pluripotency associated 2 (Dppa2) and Dppa4 are both necessary and sufficient for the activation of ZGA (De Iaco, Coudray, Duc, & Trono, 2019; Eckersley-Maslin et al., 2019). Therefore, as far as we know, at least three factors can be called master ZGA-inducer in fertilized embryos. Furthermore, overexpression of Dux is sufficient to drive pluripotent stem cells into a totipotent state by expressing 2-cell embryo specific transcripts (2-cell-like stem cells) (Macfarlan et al., 2012). In addition, several studies show totipotent state that the be achieved by depleting ZGA-repressor, such as tripartite motif-containing 28 (Trim28) (Rodriguez-Terrones et al., 2018), microRNA 34a (miR-34a) (Choi et al., 2017), chromatin assembly factor (CAF-1) (Ishiuchi et al., 2015), lysine (K)-specific demethylase 1A (Lsd1/Kdm1a) (Ancelin et al., 2016; Macfarlan et al., 2012; Wasson et al., 2016), and long interspersed nuclear element (LINE1) (Percharde et al., 2018).

The effect of ZGA-regulators in somatic cell reprograming remains unknown. Thus, the aims of this study were to explore the relationship between ZGA-regulator and SCNT efficiency, and to identify a smart factor that could promotes the efficiency of somatic reprogramming.

## Results and Discussion

We have recently established a MERVL::tdTomato based ZGA real-time monitor system (Yang et al., 2018). To identify candidate factors that facilitate ZGA in SCNT reprogramming, we carried out a siRNA-repressor and mRNA-inducer screening in SCNT (Figure 1A). From the screen, we identified a number of factors that when over-expression or knock-down resulted in reporter activation (supplement Figure S1A). Among the identified factors, Dux, Dppa2, and Dppa4 showed the strongest phenotype (Figure 1B and C). Intriguingly, we found that most SCNT embryos injected with Dppa2 or Dppa4 mRNA arrested at the 1-cell stage (called ‘1-cell block’ for simplicity; Figure 1D and E; supplement Figure S1B, Movie S1, S2). Direct injection of Dux mRNAs had no significant impact on the blastocyst formation rate (*vs.* canonical SCNT; 21.2% *vs.* 23.6%; Figure 1D; supplement Table S1). Indeed, direct injection of Dux will increase the rate of SCNT embryo fragmentation (Figure 1D and E; supplement Movie S2). Nevertheless, when Dux was over-expressed in SCNT embryos, the rate of 2-cell block was significantly reduced (Figure 1D). These results suggest that Dux (but not Dppa2/4) is a key factor whose overexpression can rescue 2-cell arrest in SCNT embryos.

**Figure 1.**
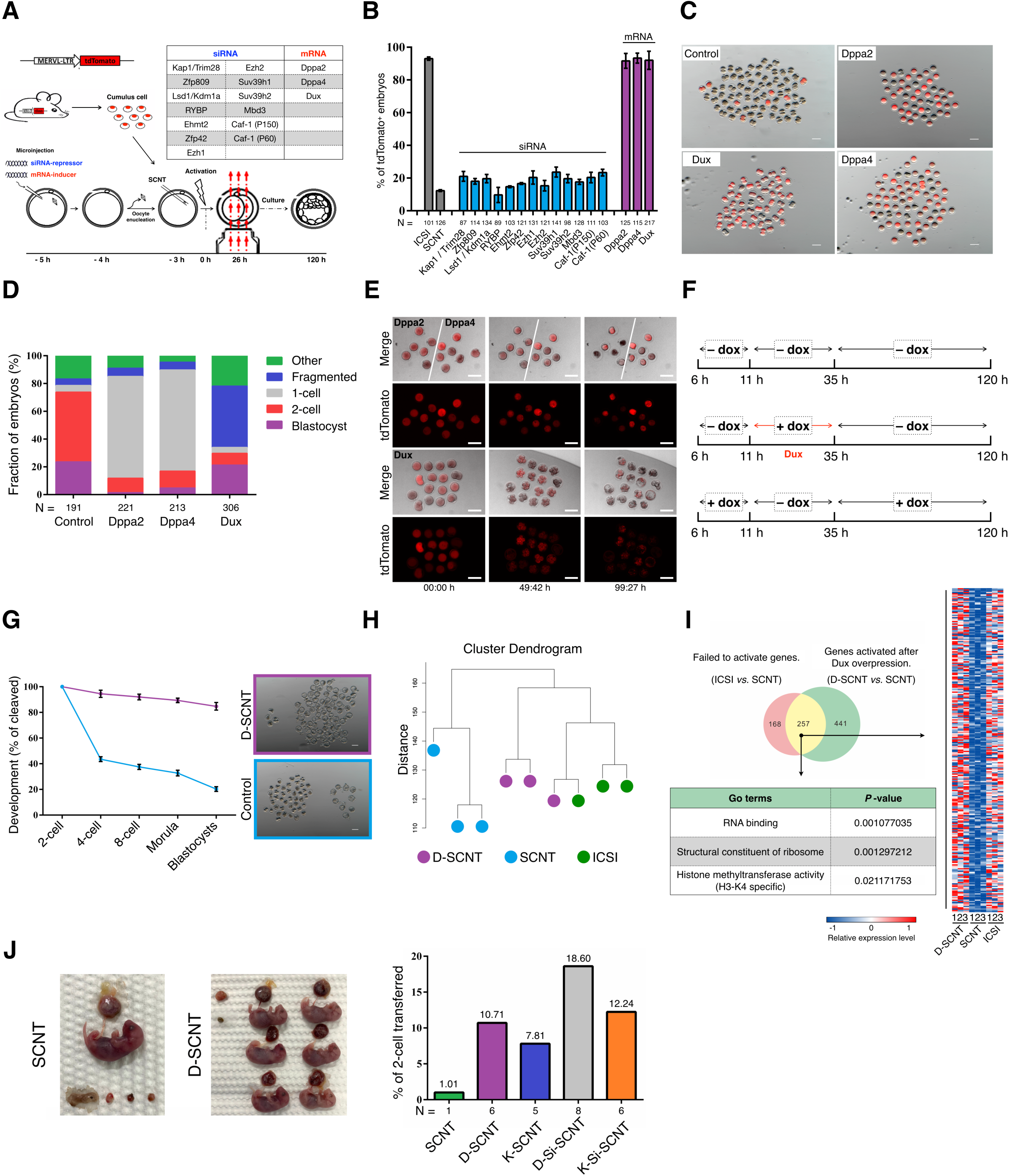
Dux act as promote factors for SCNT-mediated somatic cell reprogramming. (A) Schematic illustration of the screen strategy. (B) Quantification of embryos that expression ZGA reporter (MERVL::tdTomato) after injection with siRNA-repressor or mRNA-inducer. N, total number of embryos analyzed for each condition. Error bars, mean ± S.D.; three independent experiment replicates were performed. (C) Representative fluorescence image of SCNT embryos derived by different methods. The SCNT embryos produced by transfer of MERVL::tdTomato cumulus cell into WT enucleated oocytes. Scale bar, 100 μm. (D) Stacked bar plots show fraction of embryos at blastocyst stages after injection with different mRNA as indicated. N, total number of embryos analyzed for each condition. (E) Representative live-cell images of dynamics ZGA reporter expression during SCNT embryos development. The time after starting observation is shown on the bottom of image. Scale bar, 50 μm. (F) Schematic transient induction of Dux. All timings reported in this paper is hour post-activation. (G) The preimplantation development of SCNT embryos. Shown is the percentage of embryos that reaches each indicated stage. Scale bar, 100 μm. (H) Unsupervised hierarchical clustering. (I) Venn diagram showing the overlap between the genes that failed to be activated in SCNT 2-cell embryos and derepressed in D-SCNT. Heatmap, GO terms showing the expression pattern of 257 overlap genes. (J) The birth rate of SCNT embryos derived by different methods.

It is known that ZGA is governed by a time-dependent ‘zygotic clock’ (i.e. 24h post-fertilization), and Dux is expressed as a brief pulse in early 2-cell stage (De Iaco et al., 2019; Eckersley-Maslin et al., 2019; Rodriguez-Terrones et al., 2018). Therefore, we wonder whether Dux is time-depend on improving the efficiency of SCNT embryo development. To answer this question, we generated transgenic mouse lines containing doxycycline (dox) inducible Dux gene (supplement Figure S2A-B). By switching the culture medium between dox-containing and -lacking at various time points, the critical period for the dox-Dux requirement was mapped to a 24-h window (Figure 1F). When the dox was supplied in this window, the SCNT blastocyst formation rate was significantly increased (Figure 1G; supplement Table S2). Subsequently, we performed single-cell RNA sequencing (scRNA-seq) to examine the possible changes of the ZGA related gene expressions in Dux-overexpressed 2-cell SCNT embryos (D-SCNT). Unsupervised hierarchal clustering revealed that the ICSI and D-SCNT generated embryos in the same cluster, and ZGA-genes were upregulated in D-SCNT embryos (Figure 1H and I; supplement Figure S3A-D; Dataset S1, S2). These results indicating the improvement of Dux on SCNT embryonic development depends on its treatment time.

The SCNT blastocyst rate was improved by injecting Kdm4d mRNA (K-SCNT) (Matoba et al., 2014). We next compared the developmental potential of SCNT embryos derived by K-SCNT and D-SCNT. We found that the blastocyst formation rate of D-SCNT was similar to K-SCNT (84.72% *vs.* 86.79%; supplement Figure S4A, Table S2). Nonetheless, the cloned pup birth rate of D-SCNT was higher than K-SCNT (10.71% *vs.* 7.81%; Figure 1J; supplement Table S3). Despite D-SCNT or K-SCNT, the abnormally large placentas are still observed in those cloned pups (supplement Figure S4B, Table S3). Previous studies have indicated that knock-down DNA-methyltransferases Dnmt3a/b led to less placental abnormalities (Si-SCNT) (R. Gao et al., 2018). We next investigated whether combined D-SCNT and Si-SCNT could further improve SCNT embryonic development. Compared with canonical SCNT, combined K-SCNT and Si-SCNT improved the pup birth-rate from 1.01% to 12.24%, but less than D-SCNT combined with Si-SCNT (18.60%; Figure 1J). Genotyping analysis confirmed that all the cloned pups were generated from donor cells (supplement Figure S4C). The large placental phenotype was rescued by the D-Si-SCNT combined approach (supplement Figure S4B).

Deng et al. identified a unique embryonic 2-cell like status as the key molecular event that led to successful chemical reprogramming (CiPS) (Zhao et al., 2018). Thus, we tried to improve the CiPS by overexpression of ZGA inducer Dux (D-CiPS). The dox-inducible mouse embryonic fibroblasts (dDux-MEFs) cells were isolated from the transgenic mice. As shown in Figure 2A, we treated the dDux-MEFs with dox for various durations, and observed that Dux displayed remarkable effect at the stage-2 of CiPS. About 36 alkaline phosphatase-positive (AP^+^) colonies could be observed in D-CiPS wells (starting from 50,000 MEFs per well of 6-well plate), while only 26 AP^+^ colonies could be observed in the canonical CiPS well (Figure 2B and C). Consistently, the ZGA related genes were significantly upregulated after transient expression of Dux (Figure 2D). These D-CiPS induced cells were pluripotent as demonstrated by standard characterization procedures (Figure 2E-G). We found, based on bisulfite sequencing, that successful demethylation occurred in the promoters of Oct4 and Nanog in the D-CiPS induced cell lines (Figure 2H).

**Figure 2.**
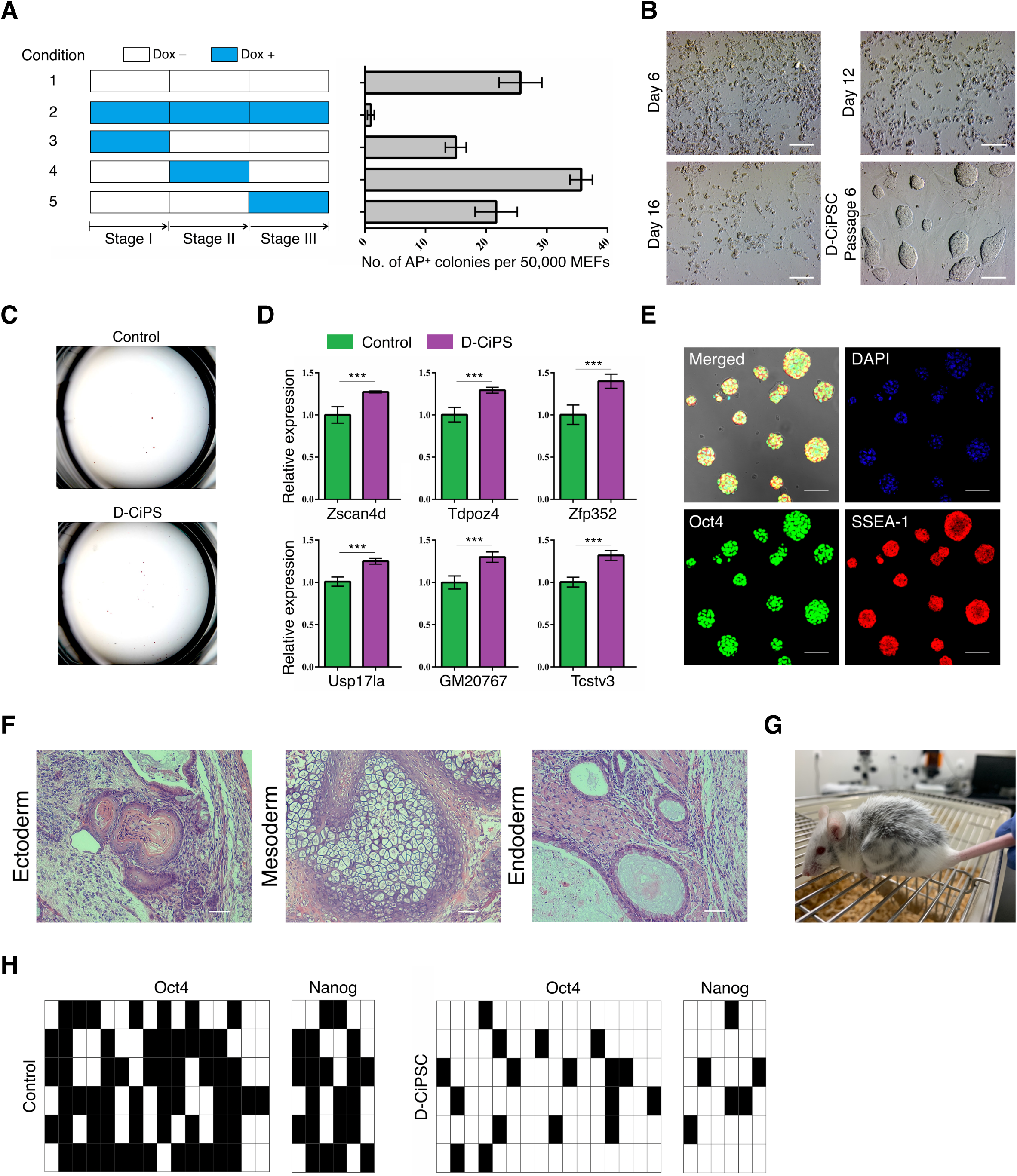
Dux facilitates chemical-mediated somatic cell reprogramming. (A) Durations of dox addition at different time points during chemical induction (left) and assessment of AP^+^ colonies (right). Error bars, mean ± S.D. (B) Morphological changes at distinct time points during D-CiPS. Scale bar, 100 μm. (C) Representative images of the number of AP^+^ positive colonies. (D) Relative expression levels by qPCR of select ZGA genes in D-CiPS, compared with canonical CiPS. Error bars, S.E.M.; n = 3; ***P < 0.001 by Student’s t-test. (E) Immunostaining images of D-CiPS expressed pluripotency markers. Scale bar, 50 μm. (F) Histology by H&E staining of teratoma tissues derived by D-CiPS cells. Scale bar, 200 μm. (G) Chimeric mouse generated from D-CiPS cells. (H) Bisulfite sequencing analysis of Oct4 and Nanog demethylation in D-CiPS or canonical CiPS cells (filled squares are methylated CpGs and empty ones indicate unmethylated).

When we prepare this manuscript, two recent studies suggested that Dux is important but not essential for fertilized embryo development by enhancing rather than ZGA inducer (Chen & Zhang, 2019; Guo et al., 2019). However, we demonstrate here that transient overexpression of ZGA-inducer Dux not only improved SCNT efficiency, but also increased the efficiency of chemical reprogramming. Ectopic expression of Dppa2/4 in SCNT embryos resulted in 1-cell block, which implied that successful timely ZGA is essential for the development of SCNT-derived embryos. A recent study found that Dppa2/4 interacted with small ubiquitin-like modifier (SUMO) related proteins (Yan et al., 2019). Therefore, we speculated that overexpression of Dppa2/4 would accelerate the degradation of reprogramming factors in the oocyte-cytoplasm, which leads to reprogramming failure. Nonetheless, fertilized embryos completely differ from SCNT-derived embryos. SCNT embryos are generated by directly injecting somatic nuclei into enucleated oocytes, whereas the oocyte cytoplasm is evolutionally designed to reprogram the spermatozoa. Therefore, although Dux and Dppa2/4 as ZGA-inducers, its effects on the reprogramming process likely differ between sperm and somatic cells. Thus, the relationship among Dux, Dppa2/4, and ZGA in SCNT warrants further investigation.

Overall, our study revealed that transient overexpression of Dux not only improved SCNT efficiency, but also increased the efficiency of chemical reprogramming. The Dux was identified as a ZGA-inducer in mouse, human, and likely all mammalian animals (Iturbide & Torres-Padilla, 2017; Whiddon et al., 2017). Thus, we reasoned that transient overexpression of Dux could also improves reprogramming efficiency of all mammals.

## Materials and methods

### Animals and Chemicals

Specific pathogen-free (SPF)-grade BDF1 (C57BL/6N × DBA/2), CD1, and Kun-Ming (KM) mice mice were purchased from Laboratory Animal Research Center (Inner Mongolia University) or Vital River Laboratories (Beijing, China). All animal experiments were approved by the Animal Care and Use Committee of Inner Mongolia University. All procedures were carried out in strict accordance with the recommendations made in the Guide for the Care and Use of Laboratory Animals of the National Veterinary and Quarantine Service. MERLV:: tdTomato transgenic mice, which carry a transgenic MERLV promoter LTR driven tdTomato reporter (Yang et al., 2018), were mated with the same positive mice. dDux-MEFs cells were isolated from E12.5 fetus. The MERLV:: tdTomato embryos used in the experiment were produced by MERVL::tdTomato sperm and MII oocytes from the littermates of transgenic mice. All chemicals used in this study were purchased from Sigma (USA), unless otherwise indicated.

### SCNT and embryo culture

The SCNT processes followed previously published studies (Kishigami et al., 2006; Yang et al., 2018). Briefly, groups of ∼ 50 MII oocytes were transferred to a chamber containing oil-covered M2 supplemented with 5 μg/ml cytochalasin B (CB). The spindle-chromosome complex (SCC) was removed by a blunt Piezo-driven pipette (∼ 10 μm internal diameter) on a 37°C heating stage of an inverted microscope (Nikon, Japan). The nuclei of donor cells were drawn in and out of the injection pipette until its plasma membrane was broken and was then injected into enucleated oocytes. The reconstructed embryos were cultured in aMEM medium (Thermo, USA) containing 10% fetal calf serum (FCS; Hyclone, USA) for 1 h before activation treatment. The reconstructed embryos were activated in Ca^2+^ free KSOM medium containing 10 mM strontium and 5 μg/ml CB for 6 h. Activated embryos were thoroughly washed and cultured in G1/G2 medium (1:1, vol / vol; Vitrolife, Sweden). Induction of Dux was performed by administration of doxycycline (2 μg/ml) in the G1/G2 medium.

### Immunofluorescence staining

Embryos and CiPS cells were rinsed three times in phosphate-buffered saline (PBS) with 0.3% BSA, fixed with 4% paraformaldehyde (PFA) overnight at 4°C, and then permeabilized with 0.2% (vol / vol) Triton X-100 for 15 min at room temperature, followed by washing thoroughly in PBS containing 0.3% BSA. Fixed samples were blocked in 0.05% Tween 20 in PBS containing 3% BSA (PBST) at 37°C for 1 h and then incubated with the primary antibodies overnight at 4°C. Samples were incubated with primary antibodies against HA (Santa Cruz, sc-7392, USA), Oct4 (Santa Cruz, sc-8629, USA), and Ssea1 (Santa Cruz, sc-21702, USA). After incubating, the samples were needed to wash several times in PBST and then incubated with appropriate secondary antibodies conjugated with Alexa Fluor 594 and Alexa Fluor 488 (Thermo, USA) for 1 h at 37°C. For imaging, the embryos were mounted in 10 μl anti-fade solution with DAPI (Thermo, USA) and compressed with a cover-slip. All samples were observed by laser scanning microscope (Nikon, Japan).

### *In vitro* mRNA synthesis and microinjection in oocytes

The *Dux, Dppa2*, and *Dppa4* mRNA were cloned into T7-driven vectors, and mRNAs were synthetized *in vitro* using mMESSAGE mMACHINE T7 Ultra Kit (Thermo, USA) following the manufacturer’s instructions. The integrity of manufactured mRNA was confirmed by electrophoresis with formaldehyde gels. The final concentration of mRNA was diluted to 900 ng/ml before injection. As previously described (Yang et al., 2018), 8 pl of mRNA was microinjected into the cytoplasm of denuded oocytes. These oocytes were obtained by superovulating BDF1 female mice by intraperitoneal injection of 10 IU pregnant mare serum gonadotropin (PMSG; Sansheng, China) and 10 IU human chorionic gonadotropin (hCG; Sansheng, China) 48 h apart. The cumulus-oocyte complexes (COCs) were collected from oviducts 14 h post-hCG, and the cumulus cells were dispersed by EmbryoMax FHM Mouse Embryo Media (Millipore, USA). Oocytes were injected using Piezo-operated blunt-end micropipette (3∼ 5 μm internal diameter). After injection, oocytes were kept at room-temperature for 30 min and then moved into the incubator. The siRNA information was presented in Table S4.

### Generation of CiPS cells

This section was adapted from Deng’s lab (Zhao et al., 2018). Reagents setup: Small molecules: VPA, CHIR99021, 616452, Tranylcypromine, Forskolin, Ch55, EPZ004777, DZNep, Decitabine, SGC0946, and PD0325901. Stage I medium: FBS/KSR-based medium supplemented with the small-molecule cocktail VC6TF5E (100 μM VPA, 40 μM CHIR99021, 10 μM 616452, 5 μM Tranylcypromine, 10 μM Forskolin, 1 μM Ch55, and 5 μM EPZ004777). N2B27-SII medium: N2B27-based medium supplemented with 10 ng/ml LIF, 50 μg/ml vitamin C, 25 ng/ml bFGF, 2 mg/ml Albumax-II, and the small-molecule cocktail VC6TF5ZDS (VPA at 1 mM). Stage III medium: N2B27-based medium with 3 μM CHIR99021, 1 μM PD0325901, 10 ng/ml LIF, and 50 μg/ml vitamin C. On day –1, MEFs were seeded at 50,000 cells per well of 6-well plate with MEF culture medium. The next day (day 0), change the medium into optimized stage I medium. From day 6∼ 12, cells were cultured in optimized N2B27-SII medium. During day 12∼ 16, cells were cultured in stage III medium with 500 μM VPA. On day 16, VPA was removed and cells were cultured in stage III medium. After another 4∼ 10 days, CiPS colonies emerged. Induction of Dux was performed by administration of doxycycline (2 μg/ml) in the culture medium.

### Transgenic mice generation

The pCW57.1-mDux-CA vector was a gift from Stephen Tapscott (Addgene 99284). The pronuclear microinjection for the production of transgenic mice followed previously published studies (Ittner & Gotz, 2007). Briefly, the vector was injected into the well-recognized pronuclei. Injected zygotes were transferred into pseudopregnant female mice (∼ 30 zygotes per mouse) after 4 h recovery culture in KSOM-AA medium (Millipore, USA). For founder identification, tail tips were subjected to standard DNA extraction procedures. The amplified DNA fragments were subjected to TA cloning and sequencing. The founder mice were crossed to the littermates of founder mice for four generations to produce homozygous Dux mice.

### Alkaline phosphatase staining

For alkaline phosphatase staining, CiPS cells were fixed with 4% PFA in PBS for 2 min, rinsed once with PBS and detection was performed using a Alkaline Phosphatase Assay Kit (VECTOR, USA) according to the manufacturer’s protocol.

### Real-time qPCR analysis

Total RNA was isolated using TRIzol reagent (Thermo, USA) and was immediately reverse-transcribed using a Prime Script RT reagent kit (Takara, Japan). The reverse transcription PCR was were amplified using Ex Taq (Takara, Japan). The qPCR was performed using a SYBR Premix Ex Taq (Takara, Japan) and signals were detected with Applied Biosystems 7500 real-time PCR System (Thermo, USA). Relative mRNA expression was calculated use the 2^(-ΔΔCt)^ method. The primer information was presented in Table S5.

### Chimeric mice and embryo transfer

For chimeric experiments, CiPS cells were used one day before passaging, which showed an optimal undifferentiated morphology. The CiPS cells were microinjected into CD1/KM blastocysts using a Piezo (Primetech, Japan) microinjection pipette. After culturing for 3 h, the embryos were transplanted into the uterus of pseudopregnant mice (∼ 15 embryos per mouse). The 2-cell stage SCNT embryos were transferred to the oviducts of E0.5 pseudopregnant (∼ 20 embryos per mouse). The cloned pups nursed by lactating CD1/KM females. SSLP analysis was performed for D6Mit15 and D2Mit102. The primer information is presented in Table S5.

### Teratoma formation

Teratoma formation analysis was carried out to evaluate the pluripotency of CiPS cells. Approximately 10^6^ CiPS cells were injected subcutaneously into the hind limbs of a 8-week-old nude mice. After 6 weeks, fully formed teratomas were dissected and fixed with PBS containing 4% PFA, then embedded in paraffin, sectioned and stained with haematoxylin and eosin for histological analysis.

### Bisulfite genomic sequencing

Genomic DNA was extracted by the tissue & blood DNA extraction kit (Qiagen, Germany) and treated with the Methylamp DNA modification kit (Epigentek, USA). The bisulphite conversion was performed as previously published studies (Cao et al., 2018).

### RNA-sequencing

The single-cell RNA-seq method followed previously published studies (W. Liu et al., 2016; Tang et al., 2010). The reverse transcription was performed directly on the cytoplasmic lysate of individual embryos. The total cDNA library was then amplified by 18∼ 20 cycles for library construction, which was performed following manufacturer’s instructions (Illumina, USA). Paired-end sequencing was further performed at Annoroad (Beijing, China). Three biological replicates were analyzed for each treatment condition. After removing low-quality reads and adapters, the raw reads were mapped to the mm9 genome using Tophat (v1.3.3) with default parameters (Trapnell, Pachter, & Salzberg, 2009). Expression levels for all RefSeq transcripts were quantified to fragments per kilobase of exon model per million mapped reads (FPKM) using Cufflinks (v1.2.0) (Trapnell et al., 2010). Gene Ontology (GO) analyses for differentially expressed genes were performed by using the R package GO.db and Database for Annotation, Visualization and Integrated Discovery (DAVID) (Huang da, Sherman, & Lempicki, 2009).

### Statistics analysis and data availability

Statistical analyses were done using the univariate analysis of variance (ANOVA) followed by the Student t-test with SPSS 21.0 statistical software (Armonk, USA). P < 0.05 was considered significant. Sequencing data have been deposited in the NCBI sequence read archive (SRA) under accession codes: SCNT-1 (SAMN12871322), SCNT-2 (SAMN12871323), SCNT-3 (SAMN12871324); D-SCNT-1 (SAMN12871325), D-SCNT-2 (SAMN12871326), D-SCNT-3 (SAMN12871327); ICSI-1 (SAMN12871328), ICSI-2 (SAMN12871329), ICSI-3 (SAMN12871330).

## Acknowledgements

We thank Prof. Shaorong Gao (Tongji University), Prof. Zhiming Han (Chinese Academy of Sciences), and Prof. Qing Xia (Peking University) for their technical assistance. We also thank Prof. Yi Zhang (Harvard Medical School) for sharing the list of RRRs. This study was supported by the Genetically Modified Organisms Breeding Major Projects (2016ZX08007-002), the opening project of State Key Laboratory of R2BGL (to L.Y.), the Inner Mongolia University Chief Scientist Program (to G.L.), the Inner Mongolia Autonomous Region Basic Research Project (to G.L.).

## Author contributions

L.Y. and G.L. conceived and designed the study. L.Y., X.L., L.S., A.D., G.S., C.B., and Z.W. performed the experiments. L.Y., X.L., L.S., A.D., and G.S. analyzed the data. G.L. supervised the project. L.Y. and G.L. wrote the paper.

