## Supplemental Figures and Tables for "Significant improvement of reprogramming efficiency by transient overexpression of ZGA inducer Dux but not Dppa2/4"

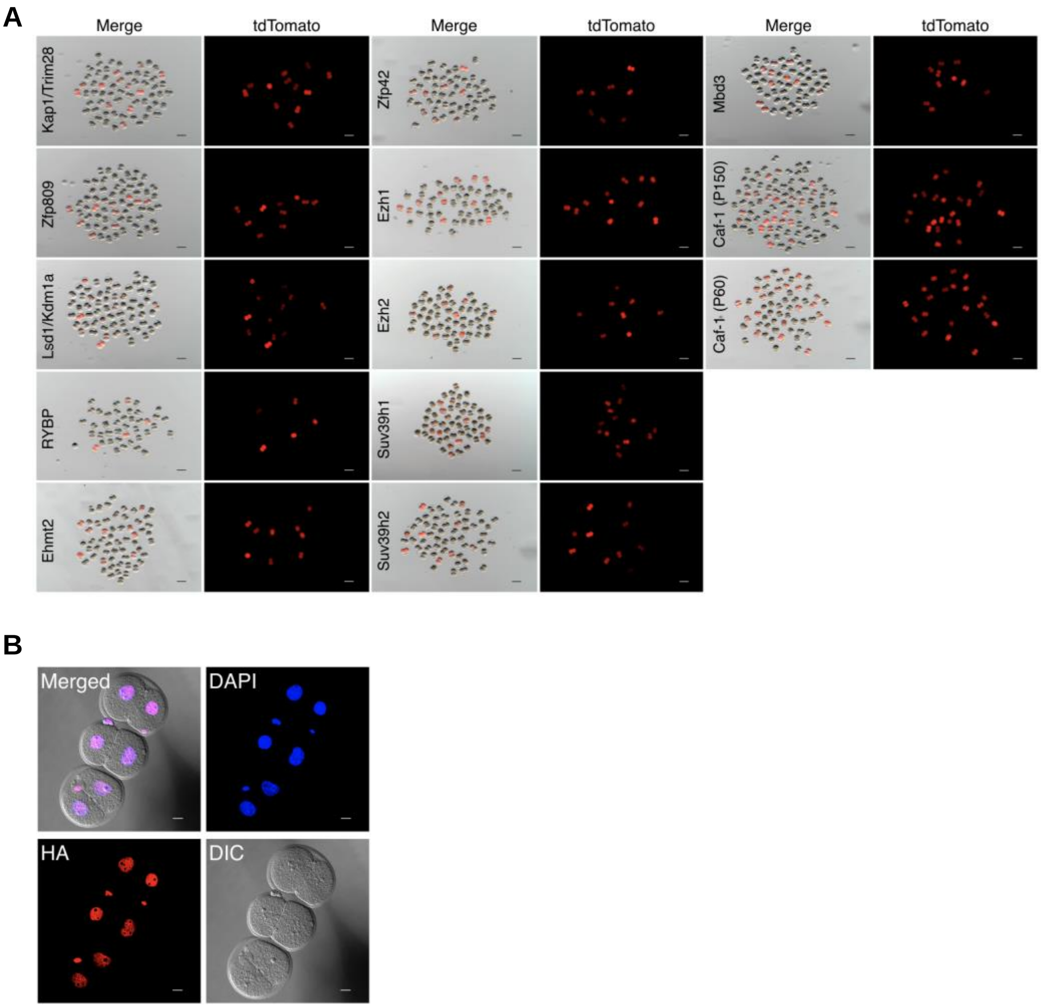


**Figure S1 Candidate genes screen and identified a list of novel ZGA regulators in SCNT embryos.**

(A) Representative fluorescence images of siRNA injected SCNT embryos. Scale bar, 100 μm.

(B) Immunostaining of SCNT embryo for HA epitope tag after injection of mRNA. Representative images from ≥ 30 embryos analyzed in three independent micromanipulations for each condition are shown. Scale bar, 20 μm.

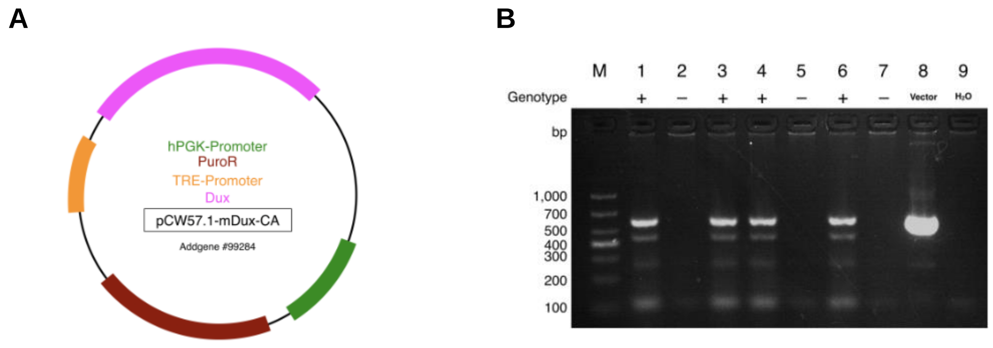


**Figure S2 Generation and characterization of dox-induced Dux transgenic mice.**

(A) Schematic representation of dox inducible Dux vector used in this study.

(B) Total DNA from transgenic founder mice (F0) and control by genomic PCR analysis. PCR products were 581 bp spanning the Dux gene. M: Marker DL1,000; lane 1, 3, 4, and 6: transgenic mice; lane 2, 5, and 7: nontransgenic mice; lane 8: positive control using pCW57.1-mDux-CA plasmid as template; lane 9: blank control using $H_2O$ as template.

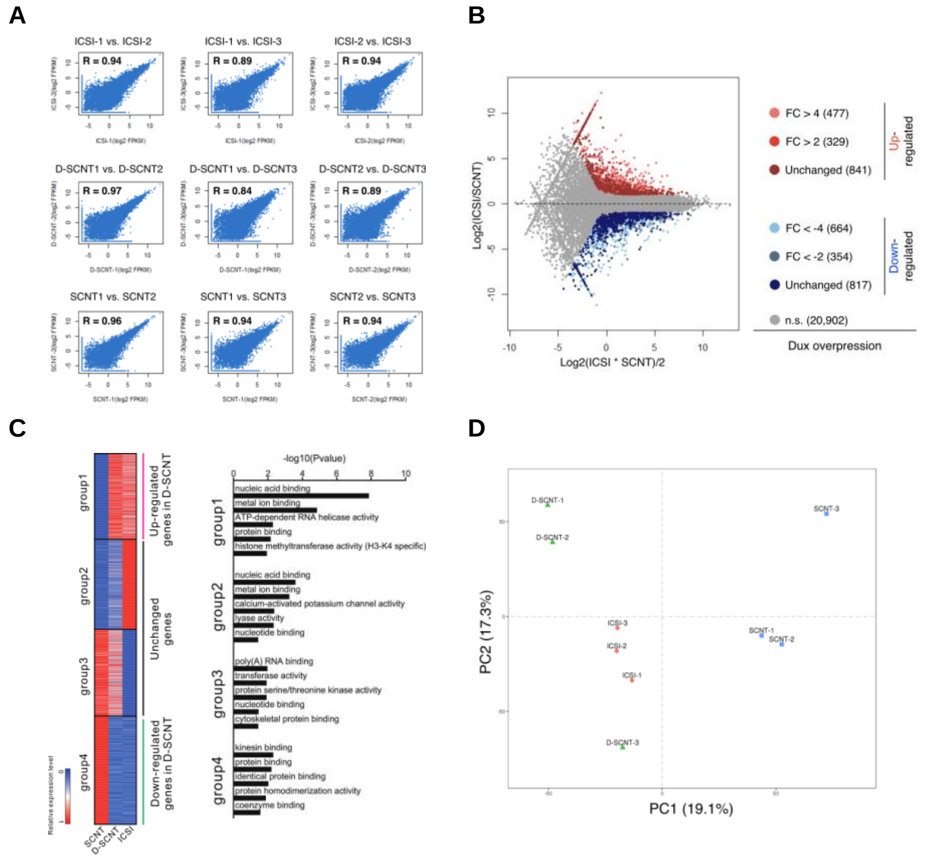

**Figure S3 Additional details on RNA-seq assay.**

(A) Scatter plot evaluation of the reproducibility of different biological replicates (Pearson correlation coefficient).

(B) The MA plot shows differentially expressed genes (DEGs) between ICSI and SCNT (all red and blue dots). The y axis represents value [log2 fold change (FC)]; x axis represents value (log2 mean expression level). Distinct color of points indicates these DEGs that are upregulated (Group 1; FC>4 or FC>2), downregulated (Group 4; FC<−4 or FC<−2) or unchanged (Group 2 and 3) in D-SCNT compared with SCNT. n.s., not significant. The total DEG genes number are shown in parentheses.

(C) Heat-map for all expressed genes patterns of DEGs among nine RNA-seq samples (FPKM > 5 in each replicate; each row represented a gene; each column represented a sample). GO analysis of the 4 groups of genes classified in Fig.S3B.

(D) PCA (principal component analysis) of RNA-seq among the ICSI, D-SCNT, and canonical SCNT embryos. The D-SCNT embryos mostly resembled the ICSI derived embryos.

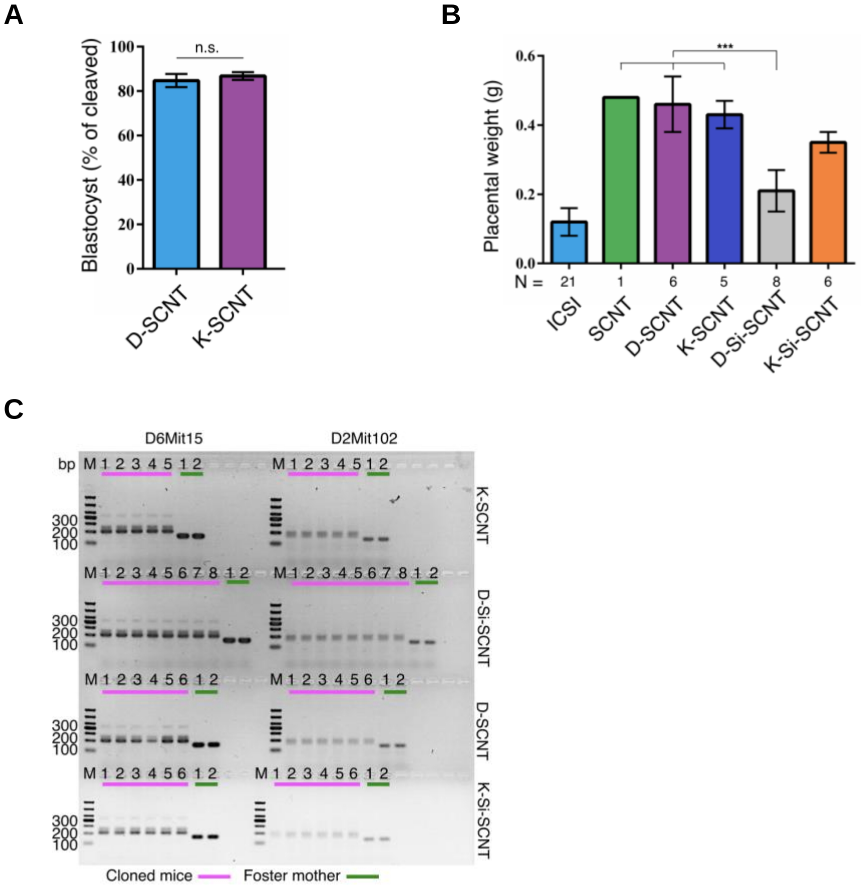


**Figure S4 Synergistic effects of D-SCNT and si3a/b-SCNT on the cloned pup birth.**

(A) The bar chart showing the efficiency of blastocyst formation. Error bars, mean ± S.D.; three independent experiment replicates were performed. n.s., not significant by Student's t-test.

(B) The bar graphs showing the placental weights of the cloned fetuses at birth. Error bars, S.D.; N, total number of placenta; ***p < 0.001 by Student's t-test.

(C) DNA genotyping identified the cloned mouse from donor cells by simple sequence length polymorphism (SSLP) analysis. M, Marker DL1,000.

**Supplement Table S1. Developmental rates of SCNT embryos injected with ZGA inducer mRNA.**

| Group | Injected with | No. of replicates | No. of cultured | No. of 2-cell (% of cultured) | No. of 4-cell (% of cultured) | No. of 8-cell (% of cultured) | No. of morula (% of cultured) | No. of blastocyst (% of cultured) |
|---|---|---|---|---|---|---|---|---|
| Control-SCNT | — | 3 | 191 | 181 (94.8) | 82 (42.9) | 76 (39.8) | 63 (33.0) | 45 (23.6) |
| Dppa2-SCNT | Dppa2 | 3 | 221 | 54 (24.4) | 27 (12.2) | 15 (6.8) | 7 (3.2) | 3 (1.4) |
| Dppa4-SCNT | Dppa4 | 3 | 213 | 51 (23.9) | 23 (10.8) | 16 (7.5) | 16 (7.5) | 10 (4.7) |
| Dux-SCNT | Dux | 4 | 306 | 292 (95.4) | 264 (86.3) | 151 (49.3) | 91 (29.7) | 65 (21.2) |

**Supplement Table S2. Developmental rates of SCNT embryos derived by canonical SCNT, K-SCNT, and D-SCNT.**

| Group | No. of replicates | No. of 1-cell embryos | % 2-cell per 1-cell ± S.D. | % 4-cell per 2-cell ± S.D. | % 8-cell per 2-cell ± S.D. | % morula per 2-cell ± S.D. | % blastocyst per 2-cell ± S.D. |
|---|---|---|---|---|---|---|---|
| SCNT | 3 | 221 | 94.53 ± 1.10 | 43.55 ±1.88 | 37.53 ± 2.03 | 32.73 ± 2.22 | 20.26 ± 1.84 |
| D-SCNT | 3 | 241 | 93.28 ± 1.20 | 94.51 ± 2.80 *** | 92.00 ± 2.22 *** | 89.40 ± 1.76 *** | 84.72 ± 2.97 *** |
| K-SCNT | 3 | 233 | 93.58 ± 1.02 | 95.39 ± 0.84 *** | 94.01 ± 0.90 *** | 90.74 ± 1.53 *** | 86.79 ± 1.71 *** |

***$P < 0.001$ as compared with SCNT group, by two-tailed Student's *t* test.

**Supplement Table S3. Postimplantation development of SCNT embryos derived by different methods.**

| Type | Treated with | No. of reconstructed 1-cell embryos | No. of 2-cell embryos (% of 1-cell) | Embryos transferred (ET) at 2-cell stage | No. of pups at birth (% of ET; normal) | Placenta weight at birth (g ± S.D.) |
|---|---|---|---|---|---|---|
| ICSI | — | 45 | 43 (95.56) | 28 | 21 (75) | 0.12 ± 0.04 *** |
| SCNT | — | 116 | 109 (93.97) | 99 | 1 (1.01) | 0.48 |
| D-SCNT | Dux | 65 | 60 (92.31) | 56 | 6 (10.71) | 0.46 ± 0.08 *** |
| K-SCNT | Kdm4b | 76 | 70 (92.11) | 64 | 5 (7.81) | 0.43 ± 0.04 *** |
| D-Si-SCNT | Dux + siDnmt3a+3b | 56 | 51 (91.07) | 43 | 8 (18.60) | 0.21 ± 0.06 |
| K-Si-SCNT | Kdm4b + siDnmt3a+3b | 58 | 54 (93.10) | 49 | 6 (12.24) | 0.35 ± 0.03 *** |

***P < 0.001 as compared with D-Si-SCNT group, by two-tailed Student's t test.

**Supplement Table S4. siRNA used in this study.**

| Gene name | Manufacturer | Catalog / PMID Number |
|---|---|---|
| Zfp809 | Santa Cruz Biotechnology | sc-155585 |
| Lsd1/Kdm1a | Santa Cruz Biotechnology | sc-60971 |
| RYBP | Santa Cruz Biotechnology | sc-77379 |
| Ehmt2 | Santa Cruz Biotechnology | sc-145298 |
| Zfp42 | Santa Cruz Biotechnology | sc-61461 |
| Ezh1 | Santa Cruz Biotechnology | sc-38188 |
| Ezh2 | Santa Cruz Biotechnology | sc-156000 |
| Suv39h1 | Santa Cruz Biotechnology | sc-38464 |
| Suv39h2 | Santa Cruz Biotechnology | sc-153944 |
| Mbd3 | Santa Cruz Biotechnology | sc-35868 |
| Kap1/Trim28 | Sequence as previously described #. | PMID 28759843 |
| Caf-1 (P150) | Sequence as previously described ##. | PMID 26237512 |
| Caf-1 (P60) | Sequence as previously described ##. | PMID 26237512 |
| Dnmt3a | Sequence as previously described ###. | PMID 30146410 |
| Dnmt3b | Sequence as previously described ###. | PMID 30146410 |

### Klimczak M, Czerwińska P, Mazurek S, et al. TRIM28 epigenetic corepressor is indispensable for stable induced pluripotent stem cell formation. Stem Cell Res. 2017;23:163–172. doi:10.1016/j.scr.2017.07.012

#### Ishiuchi T, Enriquez-Gasca R, Mizutani E, et al. Early embryonic-like cells are induced by downregulating replication-dependent chromatin assembly. Nat Struct Mol Biol. 2015;22(9):662–671. doi:10.1038/nsmb.3066

##### Gao R, Wang C, Gao Y, et al. Inhibition of Aberrant DNA Re-methylation Improves Post-implantation Development of Somatic Cell Nuclear Transfer Embryos. Cell Stem Cell. 2018;23(3):426–435. doi:10.1016/j.stem.2018.07.017

**Supplement Table S5. Primer used in this study.**

| Genes targeted | Application | Sequences (5'-3') |
|---|---|---|
| Zscan4 | Real-time PCR. | forward: AAATGCCTTATGTCTGTTCCCTATG |
| | | reverse: TGTGATAATTCCTCAGGTGACGAT |
| Tdpoz4 | Real-time PCR. | forward: ACCCAAGACCTGCAATCAAG |
| | | reverse: ATTCATGGCCAGCTACCAAC |
| Zfp352 | Real-time PCR. | forward: AAAGCCTTGATCCTCAGGTG |
| | | reverse: GCCGAAGAGTTTTTCTGAGG |
| Usp17la | Real-time PCR. | forward: TTTGTAGACACGGTGGTTGC |
| | | reverse: GGGAGCAGAAGGAAGTTTTTC |
| Gm20767 | Real-time PCR. | forward: TGCTTCCTATCCAGCTCTTG |
| | | reverse: CGGAAAAGGACTGCATCATC |
| Tcstv3 | Real-time PCR. | forward: AGAAAGGGCTGGAACTTGTGACCT |
| | | reverse: AAAGCTCTTTGAAGCCATGCCCAG |
| Gapdh | Real-time PCR. | forward: GTGGCAAAGTGGAGATTGTTG |
| | | reverse: CTCCTGGAAGATGGTGATGG |
| Dux | Genotyping of transgenic mouse. | forward: ACCCATTGGAGTTGTTTCTG |
| | | reverse: GTGCGGCTTCCGTTTGT |
| D6Mit15 | Genotyping of cloned mouse. | forward: CACTGACCCTAGCACAGCAG |
| | | reverse: TCCTGGCTTCCACAGGTACT |
| D2Mit102 | Genotyping of cloned mouse. | forward: TATTTCCCTGTCACTCCTCCC |
| | | reverse: TGTCTTTATGCTCAGACATACACA |
